# Reorganisation of Cortico-Hippocampal White Matter Pathways in Healthy Ageing: Evidence for Paradoxical Shifts in Structural Connectivity

**DOI:** 10.64898/2025.11.28.691222

**Authors:** Marshall A. Dalton, Arkiev D’Souza, Anahid Ansari Mahabadian, Fernando Calamante, Olivier Piguet

## Abstract

The hippocampus plays a central role in episodic memory and has been the focus of extensive research over past decades. A substantial body of work has demonstrated that age-related memory decline is linked to changes in how the hippocampus functionally interacts with distributed brain networks. While functional connectivity changes in ageing are well documented, relatively little is known about alterations in the structural connectivity (SC) of the hippocampus, despite its foundational role in supporting communication across neural systems. In this study, we combined high-quality data from the Human Connectome Project and advanced diffusion-weighted imaging (DWI) methods to investigate age-related changes in hippocampal SC. Using a recently developed tractography pipeline that allows greater anatomical specificity than conventional approaches, we systematically compared connectivity patterns between younger (26–30 years) and older (56–60 years) adults. Results revealed reduced hippocampal SC with the entorhinal cortex and medial parietal cortices in older participants, alongside increased SC with anterior temporal areas. This paradoxical pattern suggests that ageing is associated with both vulnerability and reorganisation of hippocampal networks, with increased hippocampal-temporal connectivity potentially reflecting compensatory plasticity in response to reduced posterior medial connection. These findings provide *in vivo* evidence of cortico-hippocampal structural reorganisation in late middle age, a critical period when pathological processes such as tau deposition are already detectable in cognitively healthy individuals. More broadly, they demonstrate the power of our anatomically refined tractography pipeline as a ‘proof of concept’ for detecting subtle, regionally specific changes in hippocampal pathway density. This approach holds promise for charting normative ageing trajectories and identifying early biomarkers of vulnerability and compensation in memory-related networks.

## Introduction

The hippocampus is central to a diverse range of cognitive and affective functions, including episodic memory (Scoville & Milner, 1957), spatial navigation (Maguire et al., 2006; Okeefe & Nadel, 1979), scene-construction, mental imagery and imagination (Dalton et al., 2018; Hassabis et al., 2007), future thinking (Addis et al., 2007), decision making (McCormick et al., 2016), and the regulation of emotion, fear, stress, and anxiety (Bartsch & Wulff, 2015). Decades of research have documented age-related alterations in hippocampal function and resting-state functional connectivity, demonstrating widespread changes in hippocampal coupling with local and distributed cortical networks (Dalton et al., 2019; Salami et al., 2014; Stark et al., 2021). In contrast, relatively little is known about how the anatomical pathways that support functional communication between the hippocampus and the cortical mantle are altered over the course of healthy ageing.

Structural connectivity (SC) underpins functional interactions across neural systems, yet the SC of the human hippocampus remains comparatively understudied. Diffusion-weighted imaging (DWI) studies have shown age-related white matter decline both globally and regionally, including within the medial temporal lobe (MTL) (Bennett & Madden, 2014; Goldstein et al., 2009). Conventional tractography approaches, however, face fundamental challenges in the hippocampus, where complex fibre crossing and partial volume effects limit anatomical specificity. As a consequence, the field has lacked the methodological precision to isolate and probe specific white matter pathways connecting the hippocampus with cortical targets and to detect subtle, regionally specific age-related changes in these pathways.

This limitation is especially salient given that the hippocampus forms direct anatomical connections with regions in which tau pathology first emerges in healthy ageing and Alzheimer’s disease, including the transentorhinal and entorhinal cortices (Braak & Braak, 1992; Lace et al., 2009). One theory proposes that tau propagates into the hippocampus along the white matter pathways that directly link it to these regions (Lace et al., 2009). The capacity to reliably isolate these discrete pathways *in vivo* using MRI, and to quantitatively characterise how they change in healthy ageing and disease is therefore critical, as these pathways represent the earliest anatomical routes through which pathology may spread and memory systems begin to destabilise. Without the tools to map specific anatomical pathways at sufficient resolution *in vivo*, the structural basis of reduced cortico-hippocampal functional connectivity in healthy ageing remains poorly understood and somewhat conjectural.

To address longstanding limitations in mapping anatomical connectivity of the *in vivo* human hippocampus, we recently developed a novel method that overcomes these constraints. This approach combines track density imaging with a customised tractography pipeline to map cortico-hippocampal connectivity with unprecedented anatomical specificity (Dalton et al., 2022). Our work in young, healthy adults has shown that this tailored approach allows biologically plausible tracking of white matter pathways into the hippocampus, incorporates manual hippocampal segmentation to improve anatomical accuracy, and leverages high-resolution track density imaging to quantitatively assess connectivity strength between the hippocampus and specific cortical targets. The result is a method capable of isolating discrete hippocampal pathways with high spatial precision in the *in vivo* human brain. To date, no study has applied this approach to investigate age-related differences in hippocampal SC.

Here, we address this crucial gap in human brain connectivity. The aims of this study were twofold: (i) to replicate previous findings of cortico-hippocampal connectivity in young healthy adults (Dalton et al., 2022) in an independent cohort, and (ii) to extend this work by applying the same approach to an older cohort in late middle age (56–60 years). This age range is of particular importance, as pathological processes associated with Alzheimer’s disease, such as tau deposition in the trans/entorhinal cortices, are already detectable in a large proportion of cognitively normal individuals in their fifties (Braak & Braak, 1997; Crary et al., 2014; Lace et al., 2009). Thus, this group provides a unique window into subtle connectivity changes that may foreshadow later cognitive decline.

We predict that, compared to the younger group, older participants will show reduced white matter connectivity between the hippocampus and the entorhinal cortex, consistent with its role as a site of early vulnerability. In addition, we conducted an exploratory analysis of hippocampal structural connectivity with the wider cortical mantle to identify other pathways that may be altered in ageing. More broadly, this work serves as a proof of concept for the utility of our recently developed anatomically precise tractography pipeline in detecting subtle age-related changes in hippocampal connectivity. In doing so, we highlight its potential as a powerful tool to quantitatively chart normative trajectories of healthy ageing and develop novel early biomarkers of pathological risk.

## Materials and Methods

All data were obtained from the Human Connectome Project - Young Adult (HCP - YA) (Van Essen et al., 2013) and Human Connectome Project - Aging (HCP - A) (Bookheimer et al., 2019) open-access databases.

### Participants

MRI imaging data from 10 younger (26-30 years; 5 female) and 10 older (56-60 years; 5 female) participants were selected from the minimally processed HCP-YA and HCP-A datasets respectively, based on scan quality and clear visibility of hippocampal outer boundaries on T1-weighted images. High-quality scans in which the entire anterior-posterior extent of the hippocampus was clearly visible were essential to ensure anatomical accuracy for manual hippocampal segmentation (described below).

### Image acquisition

For the younger group (HCP-YA), the HCP diffusion protocol was used to acquire images, consisting of three diffusion-weighted shells (b-values: 1000, 2000 and 3000 s/mm^2^, with 90 diffusion weighting directions per shell) plus 18 reference volumes (b = 0 s/mm^2^). To correct for distortion, images were acquired twice with opposite phase-encoding directions (Andersson et al., 2003). The diffusion image matrix was 145 x 145 with 174 slices and isotropic voxels of 1.25 mm. The TR and TE were 5520 and 89.5 ms, respectively. Furthermore, a high resolution T1-weighted dataset with an isotropic voxel size of 0.7 mm, TR/TE values of 2400/2.14 ms, and flip angle of 8° was included for each subject. In contrast, the ageing dataset (HCP-A) included two diffusion-weighted shells (b-values: 1500 and 3000 s/mm²), each with 92-93 diffusion-weighted volumes and 14 reference volumes (b = 0s/mm^2^).

To facilitate a closer comparison between the young and ageing datasets, the datasets were sub-sampled to match the b-values and number of directions. Specifically, for the young dataset, 14 b=0 volumes and all 90 volumes from the b=3000 s/mm² shell were selected to create a single-shell DWI image (b-values = 0 and 3000 s/mm²; 14 and 90 volumes, respectively). For the ageing dataset, 14 b=0 volumes were combined with the first 90 volumes of the b=3000 s/mm² shell to generate a comparable single-shell DWI image (b-values = 0 and 3000 s/mm²; 14 and 90 volumes, respectively).

### Manual segmentation of the hippocampus

The whole hippocampus was manually segmented on coronal T1-weighted images using ITK-SNAP (Yushkevich et al., 2006) following the manual segmentation protocol outlined by Dalton and colleagues (Dalton et al., 2017). Although the original protocol details hippocampal subfield segmentation, here we applied only the outer boundary criteria to generate whole-hippocampus masks encompassing all subfields (DG, CA4–1, subiculum, presubiculum, parasubiculum) in alignment with our previous work (Dalton et al., 2022). Care was taken to ensure no encroachment into adjacent grey or white matter structures (Figure 1).

**Figure 1.**
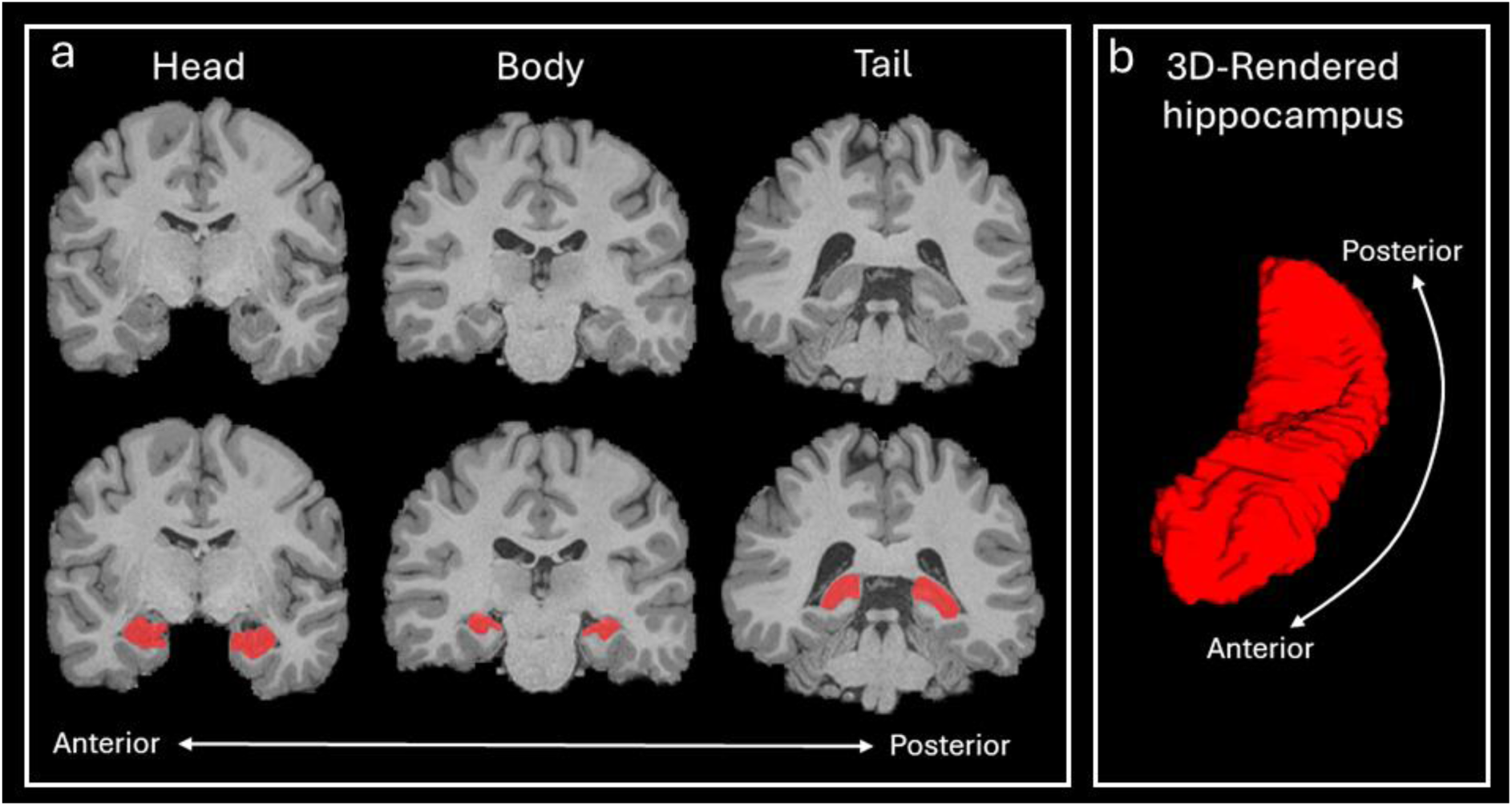
Representative example of the hippocampus mask in one participant. **(a)** Coronal slices from a T1-weighted structural image at the level of the hippocampal head (left), body (middle) and tail (right). Images in the bottom row are the same as those in the top row with the hippocampus mask overlaid (red). **(b)** A 3-D rendering of the left hippocampus mask.

### Image pre-processing and whole brain tractography

DWI data analysis was performed using the MRtrix3 software package (http://www.mrtrix.org) (Tournier et al., 2019, 2012). Full details of our preprocessing pipeline are described in Dalton and colleagues (2022). In brief, preprocessing followed guidelines from previous work (Civier et al., 2019) including bias-field correction (Tustison et al., 2010) and single-shell 3-tissue constrained spherical deconvolution to generate fibre orientation distribution (FOD) images (Jeurissen et al., 2014; Tournier et al., 2007). The T1-weighted structural image was used to generate a ‘five-tissue-type’ (5TT) image using the FSL analysis platform (Patenaude et al., 2011; Smith et al., 2012; Smith et al., 2004) to segment; (1) cortical grey matter, (2) subcortical grey matter, (3) white matter, (4) CSF and (5) ‘pathological’ tissue. In relation to tractography, the 5^th^ tissue type is considered unconstrained tissue, where no anatomical constraint is added during tracking (discussed further below). The FOD and 5TT images were used to create 70 million anatomically constrained tracks (also referred to as streamlines) across the whole brain (Smith et al., 2012) using dynamic seeding (Smith et al., 2015b) and the 2nd-order Integration over Fibre Orientation Distributions (iFOD2) probabilistic fibre-tracking algorithm (Tournier et al., 2010). These tracks are referred to as a whole brain tractogram (Figure 2a).

**Figure 2.**
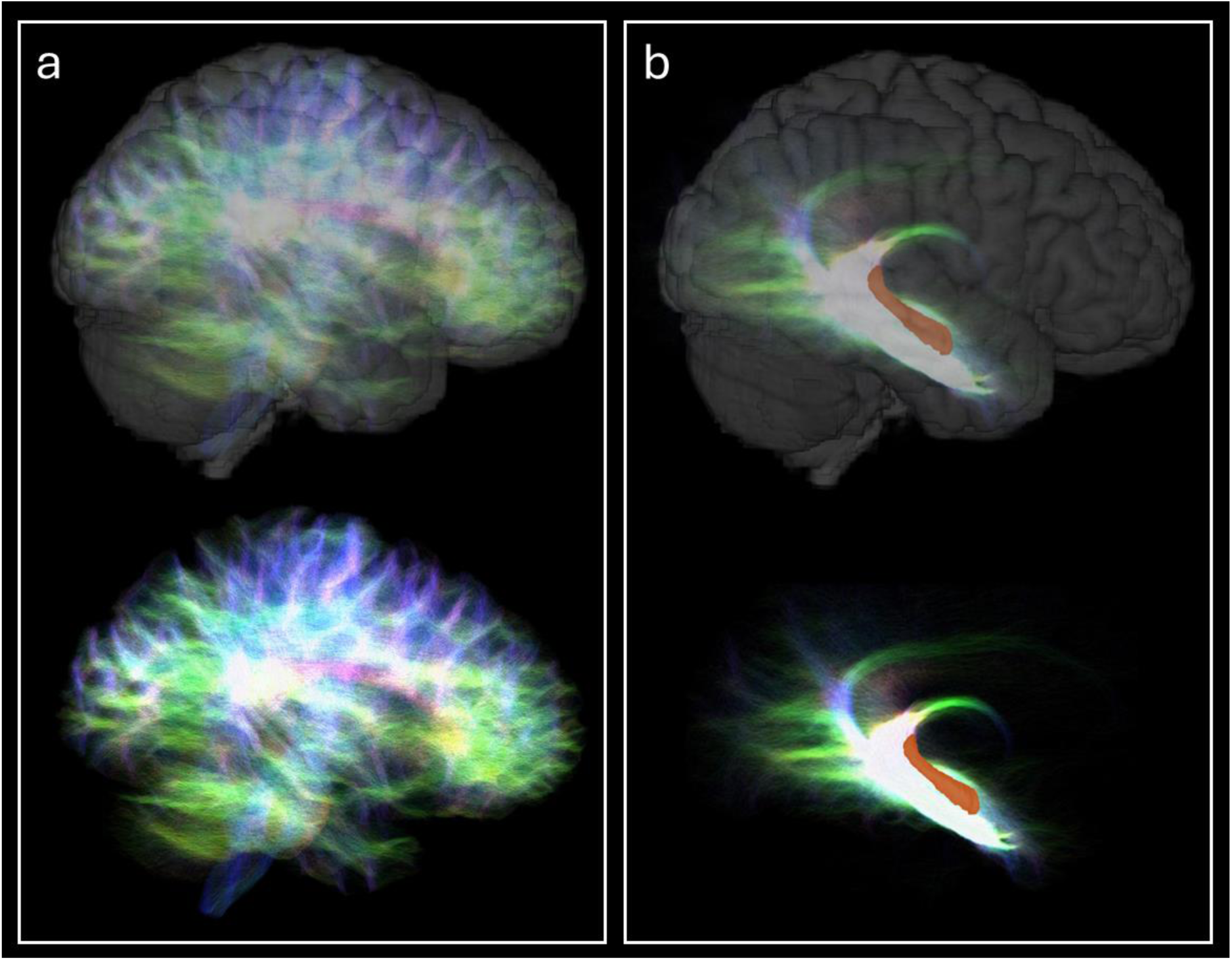
**(a)** Representative example of a whole brain tractogram (sagittal view) from a single participant and **(b)** a hippocampus tractogram from the same participant with the 3D rendered hippocampus mask overlaid (orange). Note; the hippocampus tractogram contains only the white matter pathways that terminate within the hippocampus. In each panel, the top image shows the tractogram overlaid on a 3D-rendered whole brain image and the bottom image shows the same tractogram in isolation. Tractograms are displayed with transparency; colours represent the direction of fibre orientation; high intensity represents high density of tracks.

### Hippocampus tractography

To reliably measure hippocampus specific connectivity, we applied our streamline-tracking approach (Dalton et al., 2022) which facilitates biologically plausible entry of streamlines into the hippocampus and isolates only those streamlines with an endpoint within the hippocampus. In brief, for each participant, an additional hippocampal mask was created that extends inferiorly to encompass the grey matter white matter interface (gmwmi) directly beneath the hippocampus, where white matter fibres are known to enter and exit the hippocampus (Duvernoy et al., 2005; Witter, 2007). This mask was labelled as white matter in the 5TT image, while the manually segmented whole hippocampus mask was assigned as the fifth tissue type. Note that labelling it as fifth tissue type involves not setting any constraints for tracking once the streamline is within the hippocampus. This configuration allowed streamlines to traverse the gmwmi underlying the hippocampus, where streamlines are typically terminated using conventional approaches (see Huang et al., 2021 for example) and, instead, to continue tracking within the hippocampus until streamlines reach their natural termination zones. Our approach allows streamlines to enter the hippocampus in a biologically plausible manner and ensures that only those streamlines that terminate within the hippocampus are isolated and retained for further analysis.

Using this modified 5TT image, we generated an additional 10 million streamlines seeded from the hippocampus. These streamlines were combined with the whole-brain tractogram, and spherical-deconvolution informed filtering of tractograms 2 (SIFT2) (Smith et al., 2015b) was applied to the combined 80 million-streamline file. SIFT2 allows quantitative assessment of white matter fibre density by assigning a connectivity weighting to each streamline (see Smith et al., 2015a for details). All streamlines with an endpoint in the hippocampus, along with their SIFT2 weights, were extracted to form the hippocampus tractogram (Figure 2b).

### Whole hippocampus connectivity

We applied the Human Connectome Project Multi-Modal Parcellation (HCPMMP) (Glasser et al., 2016) to each participant’s T1-weighted structural image using FreeSurfer (Fischl, 2012). This parcellation divides the cortex into 180 areas per hemisphere. We replaced the automated hippocampus mask with our manually segmented mask to ensure anatomical accuracy.

The strength of anatomical connectivity between the hippocampus and every other parcel of the HCPMMP was measured using the sum of SIFT2 weighted tracks (Smith et al., 2015b). For each cortical area, we combined the values for left and right hemisphere and report bilateral values. These values were used to assess age-related differences in the density of connections between the hippocampus and cortical areas as defined by the HCPMMP.

## Statistical Analyses

Prior to statistical testing, we observed that although distributions were generally normal, several outliers (>2.5 SD above the group mean) were detected in specific cortical areas. Single outliers were observed in 10 cortical areas: EC, V3, POS1, RSC, PHA2, V6, DVT, PHA3, Pir and V4. Importantly, these outliers did not arise from the same participants across areas, indicating that they were not attributable to systematic measurement error in a subset of individuals. Although statistical transformations could have been applied to normalise these distributions, we elected not to do so in order to avoid presenting results derived from mixed statistical approaches. Instead, we report two complementary analyses: (i) statistical tests including all values, and (ii) tests conducted after removal of outliers exceeding 2.5 SD above the group mean. For each cortical region of interest, independent-samples t-tests were performed using SPSS (Statistical Package for the Social Sciences, Version 26), and results are reported at an uncorrected significance threshold of p < 0.05.

## Results

We first characterised SC between the whole hippocampus and all cortical areas of the HCPMMP (Glasser et al., 2016). For brevity, we present results relating to the 20 cortical areas with the highest degree of SC with the whole hippocampus. Abbreviations for all cortical areas are defined in Supplementary Table 1. In the young group, the most highly connected areas included cortical areas in the medial temporal (EC, PeEC, PHA1-3), temporal (TGv, TGd, TF, Pir), parietal (POS2, POS1, RSC, DVT, ProS) and occipital (V1-4, V3A, V6) cortices (see Figure 3a and visual representation in Figure 4a). These areas are listed by rank order of connectivity strength in Table 1. These patterns broadly aligned with and replicated cortico-hippocampal connectivity patterns observed in our previous work using the same approach in a different cohort of healthy young adults (Dalton et al., 2022). In the older group, patterns of connectivity mirrored those observed in the younger group, with the same areas exhibiting the strongest connectivity with one difference. Pir in the younger group was replaced by area TE2a in the older group as one of the top 20 most highly connected areas. Overall, the global patterns of cortico-hippocampal connectivity did not differ greatly between the younger and older groups in this study.

**Figure 3.**
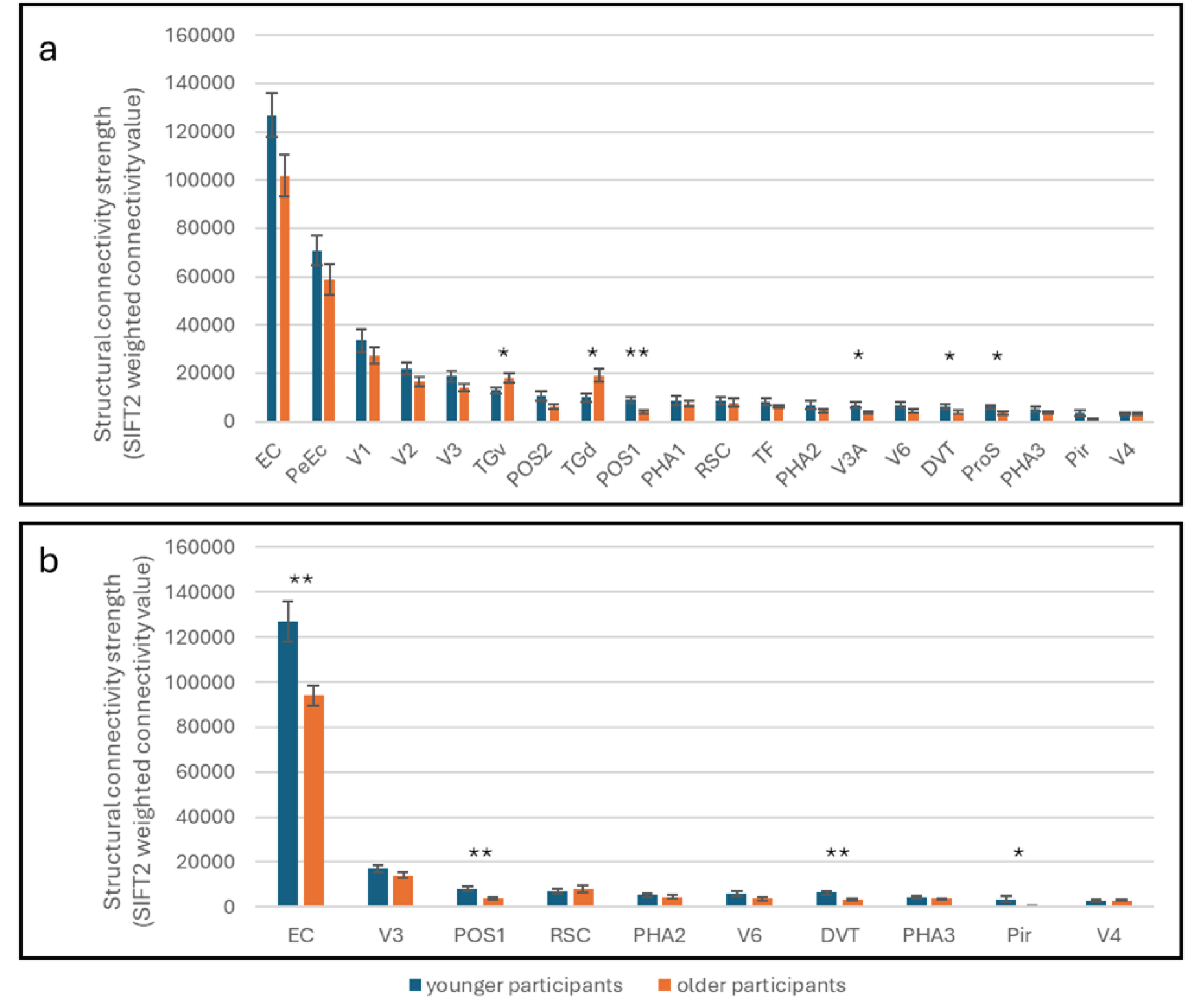
**(a)** Comparison of SC between the younger (blue) and older (orange) groups in the 20 most highly connected cortical areas. Bars represent the mean SIFT2 weighted connectivity value for each cortical area. Error bars represent the standard error of the mean. **(b)** Comparison of SC between the younger and older group for areas that contained outliers, after removal of those outliers. Bars represent the mean SIFT2 weighted connectivity value for each cortical area. Error bars represent standard error of the mean. Note, only brain areas that contained outliers are displayed in Figure 3b, as results for other brain areas are unchanged from Figure 3a. *p< 0.05, **<p0.01, uncorrected.

**Figure 4.**
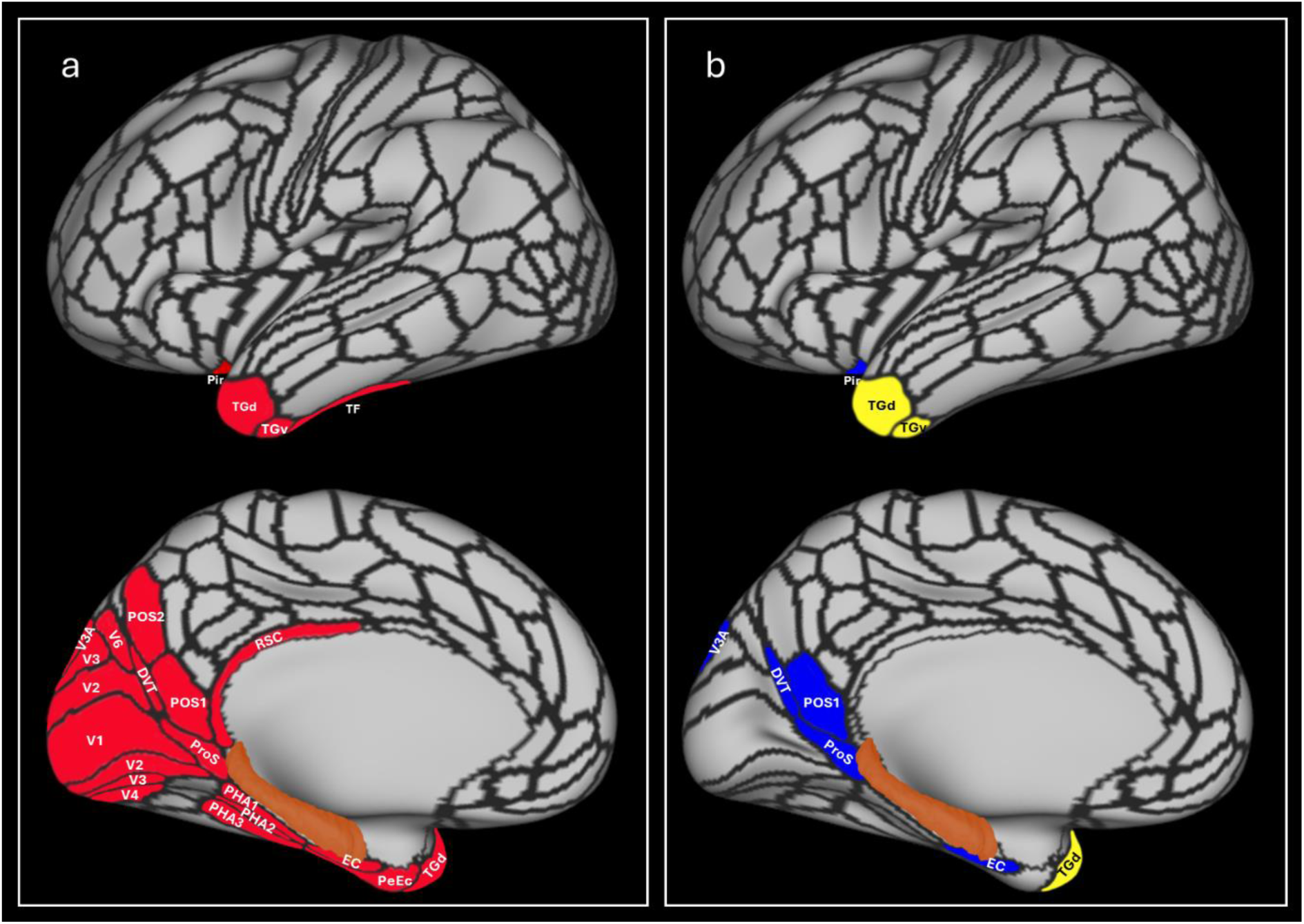
**(a)** The 20 most highly connected areas in younger participants are highlighted in red. Hippocampus is highlighted orange. **(b)** Visual representation of brain areas showing a statistically significant between group difference in connectivity strength after exclusion of outliers. Areas in blue indicate decreased connectivity in the older group and areas in yellow indicate increased connectivity in the older group. Hippocampus is highlighted orange.

**Table 1.** Statistical analysis of SC density measures of the 20 most highly connected cortical areas between the younger and older group (n.s = not significant, *p< 0.05, **p<0.01, uncorrected).

| Cortical area | Younger |  | Older |  | Independent samples t-test |  |  |
| --- | --- | --- | --- | --- | --- | --- | --- |
|  | Mean SIFT2 weighted value (connectivity strength; n=10) | Standard deviation | Mean SIFT2 weighted value (connectivity strength; n=10) | Standard deviation | t-statistic (18) | p-value | significance |
| EC | 126889 | 28220 | 101699 | 27141 | 2.035 | 0.057 | n.s |
| PeEC | 70853 | 19706 | 58929 | 20135 | 1.338 | 0.197 | n.s |
| V1 | 33498 | 14857 | 27341 | 10785 | 1.06 | 0.303 | n.s |
| V2 | 21982 | 7652 | 16593 | 5795 | 1.775 | 0.093 | n.s |
| V3 | 18741 | 6636 | 14156 | 4307 | 1.833 | 0.083 | n.s |
| TGv | 13033 | 3782 | 18134 | 6156 | -2.233 | 0.041 | * |
| POS2 | 10631 | 6045 | 6236 | 2758 | 2.091 | 0.057 | n.s |
| TGd | 9943 | 5321 | 19043 | 8755 | -2.809 | 0.013 | * |
| POS1 | 8977 | 4317 | 4052 | 1815 | 3.325 | 0.006 | ** |
| PHA1 | 8584 | 6681 | 7383 | 3499 | 0.504 | 0.621 | n.s |
| RSC | 8503 | 4952 | 7910 | 5095 | 0.264 | 0.795 | n.s |
| TF | 211 | 4503 | 6285 | 1955 | 1.24 | 0.231 | n.s |
| PHA2 | 6890 | 5829 | 4554 | 2045 | 1.196 | 0.247 | n.s |
| V3A | 6874 | 3385 | 3701 | 1098 | 2.819 | 0.017 | * |
| V6 | 6845 | 3779 | 4404 | 2880 | 1.625 | 0.122 | n.s |
| DVT | 6342 | 2855 | 3853 | 2204 | 2.182 | 0.043 | * |
| ProS | 5959 | 2221 | 3532 | 1926 | 2.612 | 0.018 | * |
| PHA3 | 5015 | 3021 | 3690 | 1384 | 1.261 | 0.223 | n.s |
| Pir | 3496 | 3956 | 830 | 970 | 2.069 | 0.065 | n.s |
| V4 | 3196 | 1916 | 3376 | 1354 | -0.243 | 0.811 | n.s |

Next, we conducted a comprehensive between-group comparison of connectivity strength by examining SIFT2-weighted density measures for the top 20 most highly connected cortical areas. For consistency, we focussed on the 20 most highly connected cortical areas in the young group, and SIFT2-weighted density values for each of these 20 areas were compared between younger and older participants.

When including all data (inclusive of outliers) in each analysis, independent-samples t-tests revealed significant between-group differences in connectivity strength between the hippocampus and several cortical areas: TGv (t(14.95) = –2.233, p = 0.041), TGd (t(14.85) = – 2.809, p = 0.013), POS1 (t(12.09) = 3.325, p = 0.006), V3A (t(10.87) = 2.819, p = 0.017), DVT (t(18) = 2.182, p = 0.043), and ProS (t(18) = 2.612, p = 0.018). In addition, trends toward significance (defined here as p < 0.06) were observed in the entorhinal cortex (EC; t(18) = 2.035, p = 0.057) and POS2 (t(12.59) = 2.091, p = 0.057). No other cortical areas showed significant between-group differences. A full set of results, including non-significant comparisons, is presented in Table 1 and Figure 3a.

In summary, relative to the younger group, the older group exhibited significantly reduced hippocampal connectivity with medial parietal and early visual cortical areas, including POS1, ProS, DVT, and V3A (Figure 4b). In contrast, the older group demonstrated significantly greater hippocampal connectivity with temporal polar areas TGv and TGd (Figure 4b).

With respect to the outliers described above, in the younger group these occurred in V3, POS1, RSC, PHA2, V6, PHA3, and V4, whereas in the older group they were observed in EC, V6, DVT, Pir, and V4.

After excluding all outliers, independent-samples t-tests revealed additional significant between-group differences in connectivity strength. Specifically, the older group showed significantly reduced hippocampal connectivity with the EC (t(13.11) = 3.29, p = 0.006) and Pir (t(9.25) = 2.336, p = 0.044) (Table 2 and Figure 3b). No other cortical areas affected by outliers exhibited significant between group differences once outliers were removed. Full statistical results for relevant areas following outlier exclusion are provided in Table 2.

**Table 2.**
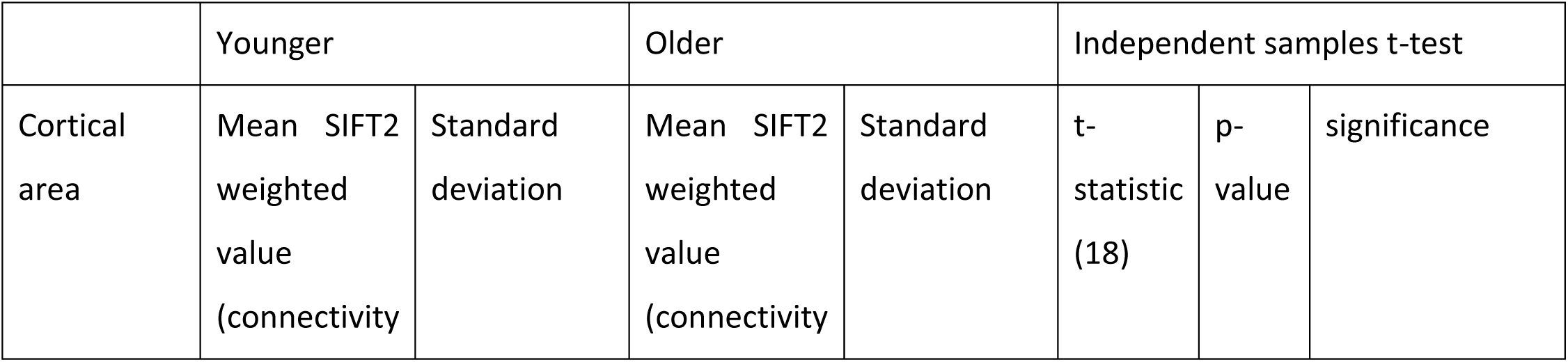
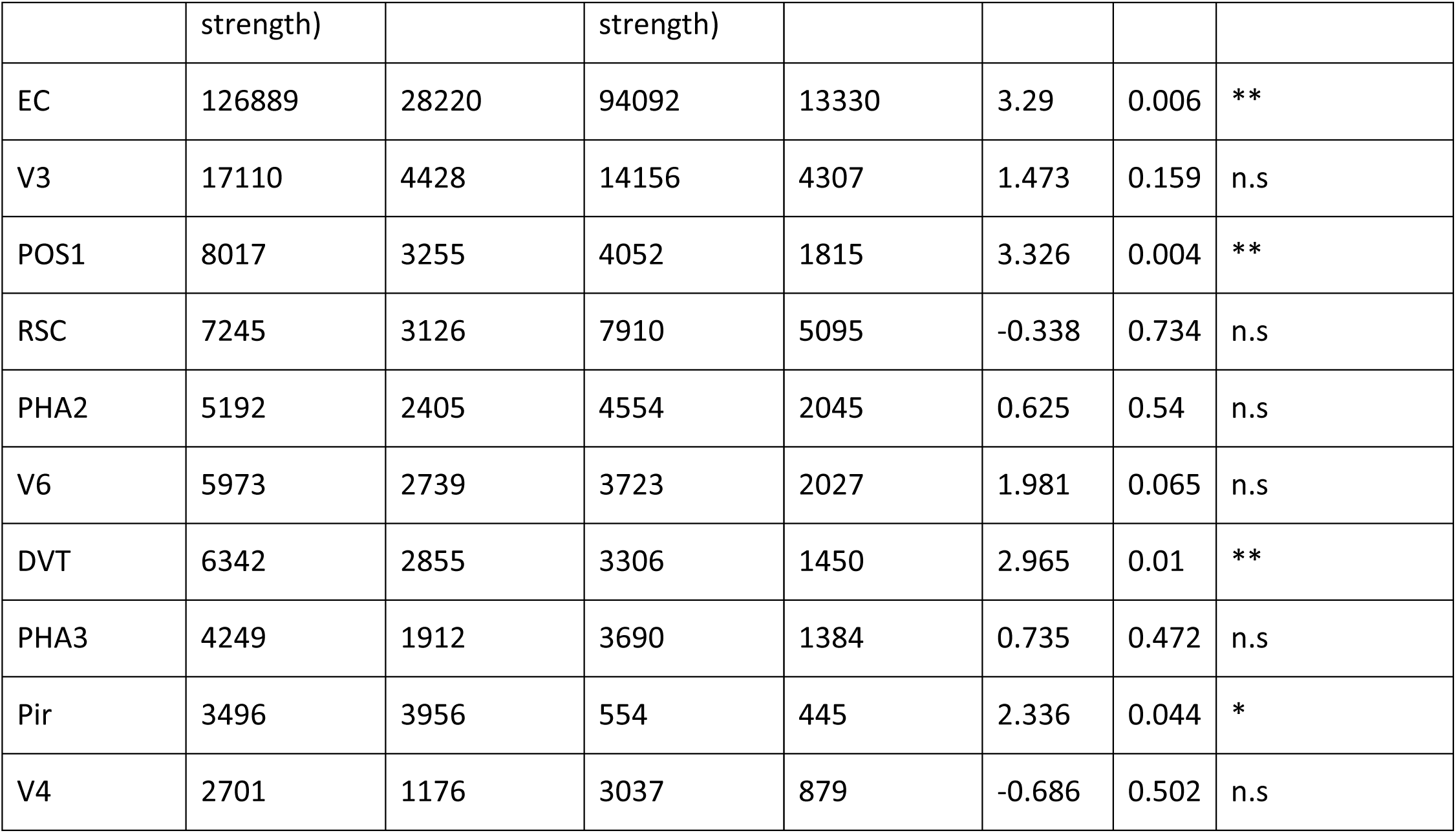
Statistical analysis of SC density measures between the younger and older groups after excluding outliers. Note, only brain areas that contained outliers are reported in this table. (n.s= not significant, *p<0.05, **p<0.01, uncorrected).

|  | Younger |  | Older |  | Independent samples t-test |  |  |
| --- | --- | --- | --- | --- | --- | --- | --- |
| Cortical area | Mean SIFT2 weighted value (connectivity) | Standard deviation | Mean SIFT2 weighted value (connectivity) | Standard deviation | t-statistic (18) | p-value | significance |

|  | strength) |  | strength) |  |  |  |  |
| --- | --- | --- | --- | --- | --- | --- | --- |
| EC | 126889 | 28220 | 94092 | 13330 | 3.29 | 0.006 | ** |
| V3 | 17110 | 4428 | 14156 | 4307 | 1.473 | 0.159 | n.s |
| POS1 | 8017 | 3255 | 4052 | 1815 | 3.326 | 0.004 | ** |
| RSC | 7245 | 3126 | 7910 | 5095 | -0.338 | 0.734 | n.s |
| PHA2 | 5192 | 2405 | 4554 | 2045 | 0.625 | 0.54 | n.s |
| V6 | 5973 | 2739 | 3723 | 2027 | 1.981 | 0.065 | n.s |
| DVT | 6342 | 2855 | 3306 | 1450 | 2.965 | 0.01 | ** |
| PHA3 | 4249 | 1912 | 3690 | 1384 | 0.735 | 0.472 | n.s |
| Pir | 3496 | 3956 | 554 | 445 | 2.336 | 0.044 | * |
| V4 | 2701 | 1176 | 3037 | 879 | -0.686 | 0.502 | n.s |

In summary, after removal of outliers, relative to the younger group, the older group exhibited additional significantly reduced hippocampal connectivity with the EC and Pir (Figure 4b).

## Discussion

In this study, we leveraged high-quality diffusion-weighted imaging data in conjunction with a recently developed tractography pipeline to investigate age-related changes in the anatomical connectivity of the human hippocampus. Overall, our findings reveal a striking reorganisation of cortico-hippocampal connectivity in the healthy ageing brain. Ageing was associated with markedly reduced hippocampal connectivity with the entorhinal cortex and key areas within the medial parietal and early visual cortices, regions central to visuospatial processing and memory-guided behaviour. In contrast, older adults showed significantly increased hippocampal connectivity with anterior temporal cortical areas, regions associated with concept-based representational systems. By quantifying connection densities between the hippocampus and the cortical mantle, we provide new evidence that healthy ageing is associated with systematic reorganisation of cortico-hippocampal connectivity. These findings provide novel insight into our understanding of healthy age-related changes in the neural architecture of the hippocampus and highlight specific pathways that may underpin both vulnerability and resilience in the ageing brain.

### Reduced hippocampal-entorhinal and hippocampal-medial parietal connectivity

A central finding of this study is an age-related decline in anatomical connectivity between the hippocampus and the EC. The EC is a key gateway for cortico-hippocampal communication, and degeneration of this pathway is thought to be a critical driver of memory decline in both normal ageing and Alzheimer’s disease (AD) (Jack et al., 2019; Maass et al., 2018). This region is among the earliest affected by tau deposition during ageing (Braak & Braak, 1992, 1995; Lace et al., 2009), and *postmortem* studies demonstrate that neurofibrillary tangles are already detectable in a majority of individuals in their late 50s (Braak & Braak, 1997; Crary et al., 2014). Although speculative, our results may provide an *in vivo* anatomical correlate of these processes in healthy late middle-age, reinforcing the notion that reduced hippocampal-EC connection is a key marker of age-related brain vulnerability. Importantly, our novel approach offers a promising pathway toward developing a biomarker capable of reliably detecting early pathological changes that are known to selectively target this circuit but are currently not reliably detectable using conventional MRI approaches.

In addition to EC changes, we observed reduced connectivity between the hippocampus and specific medial parietal cortical areas as well as early visual areas. These regions constitute a broader posterior medial network that is central to episodic memory retrieval, visuospatial processing, and scene construction (Dalton & Maguire, 2017; Dalton et al., 2018; Hassabis et al., 2007; Moscovitch et al., 2016; Ritchey et al., 2015). Consistent with this, prior functional MRI studies have reported age-related declines in hippocampal-parietal coupling (Edde et al., 2020). Our findings provide an anatomical substrate for these functional observations, demonstrating selective degradation of white matter pathways linking the hippocampus with specific medial parietal areas. Importantly, medial parietal cortices are also among the earliest sites to exhibit metabolic disruption and tau accumulation in Alzheimer’s disease (Lowe et al., 2018; Ossenkoppele et al., 2012). Our results dovetail with these functional and pathological lines of evidence and indicate that an early breakdown of hippocampal-parietal pathways in late middle-age may be a critical mechanistic link underpinning downstream age-related functional changes and the earliest stages of neurodegenerative disease.

### Increased hippocampal-temporal connectivity: A compensatory mechanism?

In contrast to the reduced connectivity observed with entorhinal and medial parietal areas, older participants displayed stronger hippocampal connectivity with anterior temporal areas (TGv and TGd) compared with younger participants. The anterior temporal cortex is closely associated with semantic memory, language, and object knowledge, which are relatively preserved in ageing compared to visuospatial memory functions (Burke et al., 1987; Chiarello et al., 1985). The observed connectivity strength shift from weakened hippocampal-parietal to strengthened hippocampal-temporal connectivity therefore provides a compelling neuroanatomical rationale for the well-documented behavioural profile of decline in episodic and visuospatial abilities alongside relative sparing of semantic knowledge in healthy ageing.

One intriguing possibility is that the increased hippocampal-temporal connectivity reflects a compensatory reorganisation of hippocampal networks in response to declining hippocampal-entorhinal and hippocampal-parietal pathway integrity. Evidence from neuroplasticity studies suggests that functional reweighting of neural circuits can preserve performance despite underlying structural decline (Barulli & Stern, 2013; Pauwels et al., 2018; Scheller et al., 2014). Future dedicated studies are needed to inform how changes in connection density observed here, may affect structure-function relationships between the hippocampus and cortical mantle. Although speculative, our findings raise the possibility that the ageing hippocampus may adaptively strengthen connections with anterior temporal regions to maintain memory function, particularly in semantic domains, when posterior medial pathways become compromised.

### Methodological considerations and limitations

While our findings provide novel insights into age-related changes in cortico-hippocampal connectivity, several limitations warrant consideration. Our findings should be interpreted with caution in view of the uncorrected results reported here. The sample size was modest (10 participants per group), primarily reflecting the time-intensive nature of manual hippocampal segmentation and creation of additional masks that extend inferiorly to cover the underlying gmwmi, a key component of our novel pipeline. Taken together, the results presented here represent a proof of concept that our novel diffusion pipeline is able to detect subtle changes in specific cortico-hippocampal connectivity pathways, even in a comparatively understudied middle-age (56-60 yo) age bracket. Future work will extend these preliminary findings to investigate a more comprehensive lifespan cohort, that will enable a more detailed assessment of these connectivity changes in a larger and more representative sample.

The HCP-YA and HCP-A groups have different DWI acquisition protocols, although we sought to match them as closely as possible in terms of number of directions and b-values. As a result, minor protocol-related differences may still influence the findings. However, despite using a modified HCP-YA DWI dataset in the current study, our results replicate those previously reported using unmodified HCP-YA data (Dalton et al., 2022). Importantly, our replication of these previously reported connectivity patterns in an independent young cohort provides further confidence in the validity of our approach. Results of our current study offer additional assurance in its utility for detecting subtle cortico-hippocampal white matter pathway density changes that may be missed using conventional approaches.

Diffusion MRI tractography also remains an indirect measure of white matter connectivity and is constrained by image resolution and modelling assumptions. Our tailored approach, however, helped circumvent these constraints in several key ways. First, we manually amended the gmwmi inferior to the hippocampus, a region where white matter pathways are known to enter and exit the structure. This adjustment allowed streamlines to permeate the hippocampus in a biologically plausible manner, avoiding the common problem of premature termination at the gmwmi. Second, by incorporating a customised hippocampus mask as a fifth tissue type, we enabled streamlines to move within the hippocampus and terminate naturally, thereby overcoming modelling assumptions that otherwise force potentially implausible terminations. Third, by combining these anatomical constraints with state-of-the-art tractography methods (ACT, iFOD2) and streamline weighting (SIFT2), we achieved quantitative connectivity estimates that more faithfully reflect underlying axonal density. Finally, by leveraging track-density imaging at high spatial resolution, our pipeline localised the distribution of streamline endpoints within the hippocampus with unprecedented detail, allowing us to reliably isolate streamlines with an endpoint in the hippocampus with greater anatomical specificity.

### Conclusion and implications

This study provides, to our knowledge, the first *in vivo* demonstration of age-related reorganisation in cortico-hippocampal structural connectivity using quantitative track density imaging combined with our tailored hippocampal tractography approach. The identification of reduced hippocampal connectivity with the EC and medial parietal cortices, alongside increased connectivity with anterior temporal regions, points to a potential shift from visuospatial-based to concept-based representational systems in healthy ageing. These findings have important implications for basic neuroscience, offering mechanistic insight into age-related memory changes, and for clinical neuroscience, suggesting candidate pathways that may serve as early biomarkers of vulnerability and compensation.

Extending this work to broader age ranges and longitudinal designs will be crucial for elucidating whether increased hippocampal-temporal connectivity reflects adaptive compensation or paradoxically indicates a trajectory toward network-level decline. Moreover, a lifespan perspective will help establish normative trajectories of cortico-hippocampal connectivity, distinguishing patterns of healthy reorganisation from those that may signal pathological change. By identifying specific pathways that weaken (hippocampal-entorhinal and hippocampal-parietal) and others that may strengthen in compensation (hippocampal-temporal), our tailored method captures distinct patterns of hippocampal connectivity change associated with late middle-age. This approach has potential to form the basis of novel imaging biomarkers capable of detecting vulnerability decades before cognitive symptoms emerge. Ultimately, our methodology and findings open the door to targeted investigations that link connectivity shifts to behavioural outcomes, and to translational work aimed at improving early diagnosis, monitoring disease progression, and tailoring interventions to support resilience in the ageing brain.

## Acknowledgements

Data were provided by the Human Connectome Project, WU-Minn Consortium (principal investigators: David Van Essen and Kamil Ugurbil; 1U54MH091657) funded by the 16 NIH Institutes and Centers that support the NIH Blueprint for Neuroscience Research and by the McDonnell Center for Systems Neuroscience at Washington University, St. Louis, MO. Additional data were provided by the Human Connectome Project Aging (HCP-A), supported by the National Institute on Aging (NIA) and the National Institute of Mental Health (NIMH) of the National Institutes of Health. We are grateful for the support of the Australian Research Council (grant numbers DP240102161 and DP210102378). OP is supported in part by a National Health and Medical Research Council of Australia Leadership Fellowship (GNT2008020). The authors acknowledge the technical assistance provided by the Sydney Informatics Hub and Sydney Imaging, two Core Research Facilities of the University of Sydney, Australia and the facilities and scientific and technical assistance of the National Imaging Facility, a National Collaborative Research Infrastructure Strategy (NCRIS) capability, at Sydney Imaging, the University of Sydney.

---

**Supplementary Table 1.**
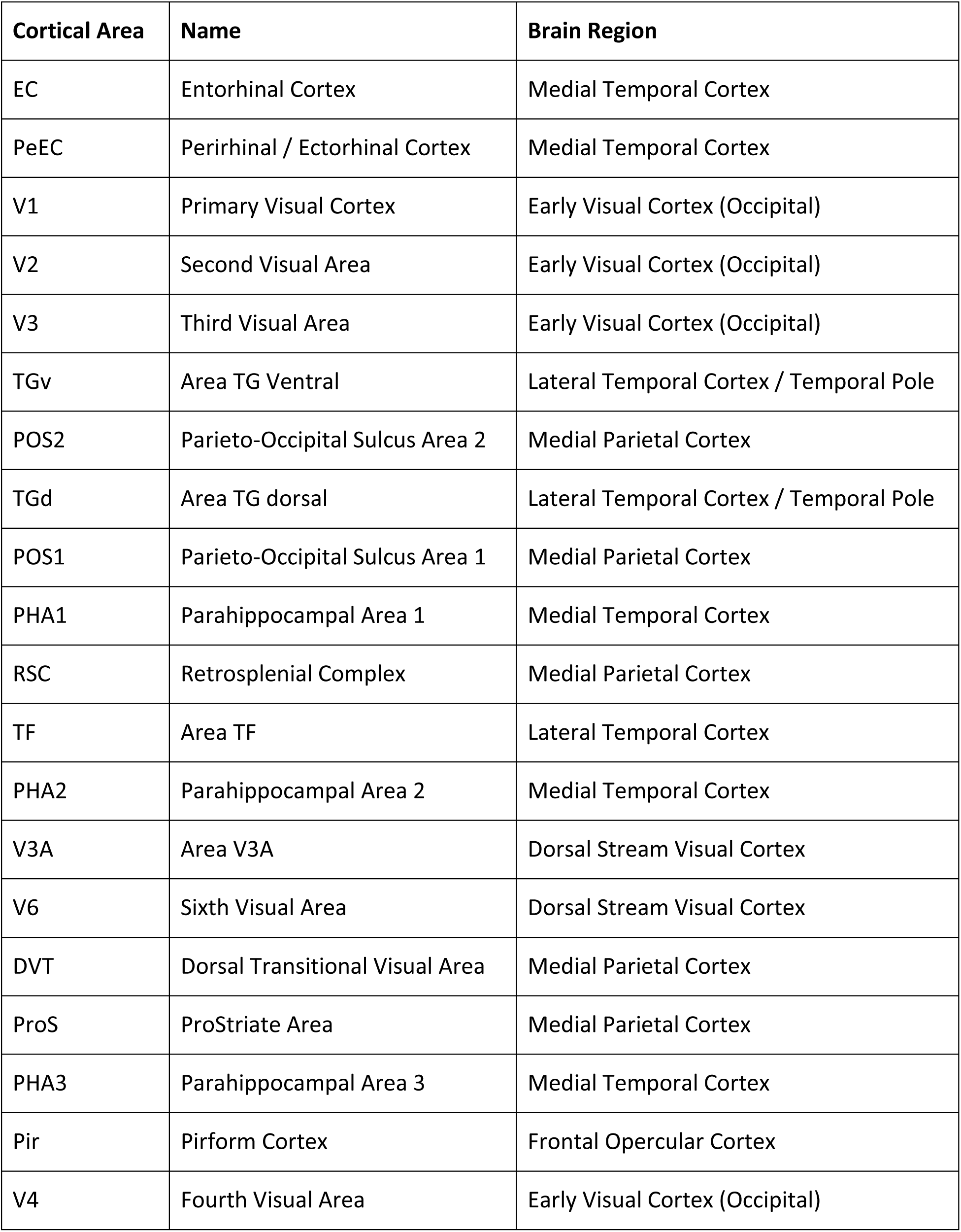
List of abbreviations for the Top 20 most highly connected cortical areas. Listed by rank order of connectivity strength in the younger group.

