## Supplementary Table 1. for "Reorganisation of Cortico-Hippocampal White Matter Pathways in Healthy Ageing: Evidence for Paradoxical Shifts in Structural Connectivity"

**Supplementary Table 1.** List of abbreviations for the Top 20 most highly connected cortical areas. Listed by rank order of connectivity strength in the younger group.

| **Cortical Area** | **Name** | **Brain Region** |
|---|---|---|
| EC | Entorhinal Cortex | Medial Temporal Cortex |
| PeEC | Perirhinal / Ectorhinal Cortex | Medial Temporal Cortex |
| V1 | Primary Visual Cortex | Early Visual Cortex (Occipital) |
| V2 | Second Visual Area | Early Visual Cortex (Occipital) |
| V3 | Third Visual Area | Early Visual Cortex (Occipital) |
| TGv | Area TG Ventral | Lateral Temporal Cortex / Temporal Pole |
| POS2 | Parieto-Occipital Sulcus Area 2 | Medial Parietal Cortex |
| TGd | Area TG dorsal | Lateral Temporal Cortex / Temporal Pole |
| POS1 | Parieto-Occipital Sulcus Area 1 | Medial Parietal Cortex |
| PHA1 | Parahippocampal Area 1 | Medial Temporal Cortex |
| RSC | Retrosplenial Complex | Medial Parietal Cortex |
| TF | Area TF | Lateral Temporal Cortex |
| PHA2 | Parahippocampal Area 2 | Medial Temporal Cortex |
| V3A | Area V3A | Dorsal Stream Visual Cortex |
| V6 | Sixth Visual Area | Dorsal Stream Visual Cortex |
| DVT | Dorsal Transitional Visual Area | Medial Parietal Cortex |
| ProS | ProStriate Area | Medial Parietal Cortex |
| PHA3 | Parahippocampal Area 3 | Medial Temporal Cortex |
| Pir | Pirform Cortex | Frontal Opercular Cortex |
| V4 | Fourth Visual Area | Early Visual Cortex (Occipital) |
